# Parallel inference of hierarchical latent dynamics in two-photon calcium imaging of neuronal populations

**DOI:** 10.1101/2021.03.05.434105

**Authors:** Luke Y. Prince, Shahab Bakhtiari, Colleen J. Gillon, Blake A. Richards

## Abstract

Dynamic latent variable modelling has provided a powerful tool for understanding how populations of neurons compute. For spiking data, such latent variable modelling can treat the data as a set of point-processes, due to the fact that spiking dynamics occur on a much faster timescale than the computational dynamics being inferred. In contrast, for other experimental techniques, the slow dynamics governing the observed data are similar in timescale to the computational dynamics that researchers want to infer. An example of this is in calcium imaging data, where calcium dynamics can have timescales on the order of hundreds of milliseconds. As such, the successful application of dynamic latent variable modelling to modalities like calcium imaging data will rest on the ability to disentangle the deeper- and shallower-level dynamical systems’ contributions to the data. To-date, no techniques have been developed to directly achieve this. Here we solve this problem by extending recent advances using sequential variational autoencoders for dynamic latent variable modelling of neural data. Our system VaLPACa (Variational Ladders for Parallel Autoencoding of Calcium imaging data) solves the problem of disentangling deeper- and shallower-level dynamics by incorporating a ladder architecture that can infer a hierarchy of dynamical systems. Using some built-in inductive biases for calcium dynamics, we show that we can disentangle calcium flux from the underlying dynamics of neural computation. First, we demonstrate with synthetic calcium data that we can correctly disentangle an underlying Lorenz attractor from calcium dynamics. Next, we show that we can infer appropriate rotational dynamics in spiking data from macaque motor cortex after it has been converted into calcium fluorescence data via a calcium dynamics model. Finally, we show that our method applied to real calcium imaging data from primary visual cortex in mice allows us to infer latent factors that carry salient sensory information about unexpected stimuli. These results demonstrate that variational ladder autoencoders are a promising approach for inferring hierarchical dynamics in experimental settings where the measured variable has its own slow dynamics, such as calcium imaging data. Our new, open-source tool thereby provides the neuroscience community with the ability to apply dynamic latent variable modelling to a wider array of data modalities.

## INTRODUCTION

Dynamic latent variable modelling has been a hugely successful approach to understanding the function of neural circuits. For example, it has been used to uncover previously unknown mechanisms for computation in the motor cortex ^1,2^, somatosensory cortex ^3^, and hippocampus ^4^. However, the success of this approach is largely limited to datasets where the observed variables have dynamics whose timescales are much faster than the dynamics of the underlying computations. This is the case, for example, with spiking data, where the dynamics governing the generation of individual spikes are much faster than the dynamics of computation across the circuit. This makes it possible to characterise the observed data, e.g. the spiking data, as a set of point-processes that can be used directly for inferring latent variables.

However, many datasets in the life sciences are generated by a hierarchy of dynamical systems, wherein the shallower-level dynamical systems that directly generate the observed data have temporal dynamics whose timescales overlap with that of the deeper-level dynamical system to be inferred. A clear example of this is *in-vivo* calcium imaging data, which is widely used in neuroscience. Thanks to advances in imaging technology and genetically encoded calcium indicators, calcium imaging enables monitoring of the activity of large populations of genetically targeted neurons in awake behaving animals ^5,6^. However, calcium imaging introduces an additional layer of a relatively slow dynamical system between the computations occurring in the brain and the measurements that neuroscientists make. This problem is outlined in Figure 1A, in which calcium fluorescence observations, *x*, depend on the state of calcium flux, *z*_1_, which is governed by a shallower-level dynamical system with temporal dynamics on the order of hundreds of milliseconds. These dynamics are driven, in part, by perturbations due to spikes, *u*_1_, which are themselves governed by a computational state, *z*_2_, with a similar timescale in its dynamics to the calcium flux (and which itself can be perturbed independently by unknown inputs, *u*_2_, that may also have slow dynamics). Due to the overlap in timescales, it is impossible to identify immediately which components of the calcium fluorescence data are driven by the dynamics of calcium flux, and which are driven by the deeper-level latent dynamics of neural computation. Ideally, neuroscientists would have a method for inferring both the shallower-level calcium dynamics and the deeper-level computational dynamics in order to uncover the hierarchical dynamical systems that generated their data. Such a tool would significantly benefit the systems neuroscience community.

**Figure 1:**
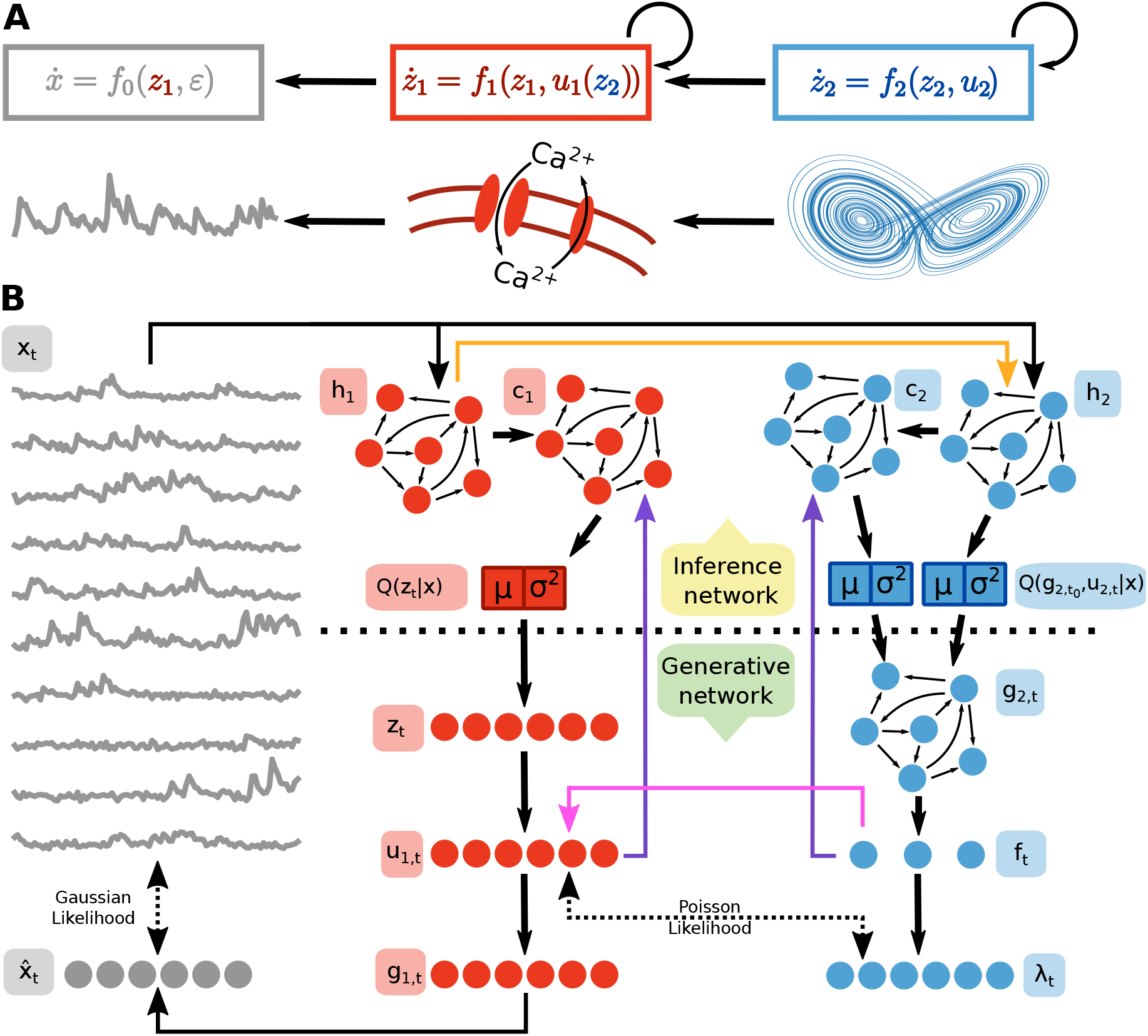
A) Hierarchy of dynamical systems (Top). schema of calcium and Lorenz dynamics (Bottom). B) Schema of our hierarchical model. Latent dynamics model in blue (right column), calcium dynamics model in red (middle column).

Currently, these problems are treated separately. For situations where the observed data can be treated as a point process, we have good techniques for inferring the deeper-level dynamics. For example, recent applications of sequential variational autoencoders with recurrent neural networks have seen great success in inferring underlying computations from extracellular spiking data ^2^. This technique, named Latent Factor Analysis of Dynamical Systems (LFADS), has improved neuroscientists’ ability to infer underlying neural computations from spiking data, e.g. it has been used to identify latent rotational reaching dynamics and to decode reaching behaviour of macaques and humans with higher fidelity than other techniques. Many other approaches using deep neural networks have also been successful in finding latent temporal structure in population spiking data, including those using piece-wise linear recurrent neural networks ^7^, generative adversarial networks ^8^, transformers ^9^, and self-supervised dual-predictor networks ^10^.

Although LFADS has significantly advanced our ability to analyze neural data in the form of spike trains, it does not address the problem highlighted above for calcium imaging, wherein the calcium dynamics introduce an additional shallower-level dynamical system whose timescale overlaps with the timescale of neural computation. In a calcium imaging scenario, what we observe is a *filtered* point process with emission noise, also known as a shot-noise process, in which our observations can be modelled as a point process that triggers an event with a stereotyped dynamical profile, along with independent white noise. In reality, this is also what we observe in electrophysiological experiments. However, the dynamics of events in such cases do not distort the inference of point-processes to the same degree.

Theoretically, this problem could be solved independently by first inferring spikes from calcium data, whether by deconvolution (e.g., OASIS) ^11^, variational inference (e.g., DeepSpike) ^12^, dynamic programming ^13^, or multiple other methods ^14–17^, and then applying LFADS. However, this approach treats each calcium trace as a completely independent variable when inferring calcium dynamics. This ignores correlations in population activity that inform the separation of calcium dynamics (which are independent of population activity) from computational dynamics (which are not independent of population activity). If this separation is sub-optimal, then inference of the deeper-level system will be impaired.

Here, we address this problem by extending LFADS with a variational ladder autoencoder architecture ^18^ that folds the calcium dynamics inference into the larger inference problem (Fig. 1B). Our system, VaLPACa (Variational Ladders for Parallel Autoencoding of Calcium imaging data), incorporates inductive biases for calcium dynamics and, thanks to the ladder architecture, is able to infer the deeper-level dynamical system better than an approach that treats inference of calcium dynamics and deeper-level dynamics as separate problems. Hierarchical Bayesian approaches to extracting neural population level statistics from calcium imaging data have had success in past, e.g. in estimating connectivity in neuronal networks ^19^ and detection of discrete activity motifs in neuronal assemblies ^20^, but none have yet directly attempted to infer latent low-dimensional nonlinear dynamics in population activity.

First, we show, using synthetic data, that we are able to reconstruct ground-truth latent dynamics from synthetic calcium traces. Next, we apply VaLPACa to spiking data from macaque motor cortex that has been converted into calcium fluorescence traces using a calcium dynamics model. We show that we are able to recover rotational dynamics from this “calcified” data just as LFADS identifies rotational dynamics from spiking data. Finally, we show using real 2-photon calcium imaging data from mouse primary visual cortex that VaLPACa can identify deeper-level latent factors that carry information about unexpected visual stimuli. Altogether, our work shows the benefits of incorporating the calcium dynamics inference procedure into the larger inference problem. It also provides neuroscientists with a new, open-source tool for analyzing calcium imaging data in order to identify deeper-level dynamics. Given the importance of calcium imaging to modern systems neuroscience, we believe that VaLPACa will be a very useful analysis tool for the community. Furthermore, our VaLPACa could be adapted to other neuroscience data modalities, such as fMRI data which, like calcium imaging, comprises both fast deeper-level latent brain dynamics and slower shallower-level blood oxygen measurement dynamics ^21^. Beyond this, we believe that our approach may be applicable to other life sciences domains, for example insurance claim modelling, where better identifying the relatively slow dynamics underlying the observed variables, such as claim submission distributions over time, could improve forecasting accuracy ^22^.

## MODEL DEFINITION

VaLPACa extends the variational ladder autoencoder (VLAE) approach ^18^ with recurrent neural networks (RNNs) (Fig 1B, Fig S1 for full directed acyclic graph), which enables inference of hierarchical latent dynamical systems. VLAEs were proposed as a way to prevent hierarchical variational autoencoders loading all of the latent code onto the shallowest level of the hierarchy (the problem of latent variable collapse in hierarchical inference). They do this by positioning more expressive networks deeper in the hierarchy to ensure that that deeper- and shallower-level features could be stored in different parts of the latent state. Crucically, no hierarchy is assumed among stochastic latent variables directly.

More explicitly, For layers 1: *L* in the hierarchy

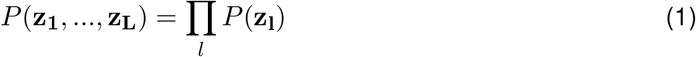

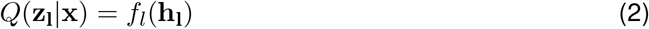

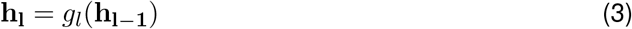

where **h**_**0**_ = **x**, and *g*_*l*_ and *f*_*l*_ are neural networks with increasing degrees of expressivity as *l* increases. This encourages more abstract features to be represented in certain portions of the latent state.

This is in contrast to traditional hierarchical autoencoders that assume a Markovian structure in the latent variables

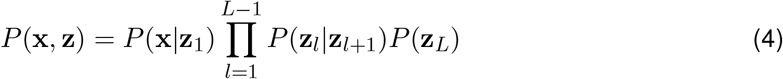

The key insight from ^18^ is that under idealised assumptions for sampling and maximization of the ELBO, the ELBO for hierarchical VAEs in this form can be optimised by minimizing the KL divergence for the shallowest layer of the model hierarchy *D*_KL_(*Q*(**z**_1_|**x**)||*P* (**z**_1_|**x**)). This means that traditional hierarchical VAEs will tend to load all representations onto the shallowest level of the hierarchy.

In contrast, by splitting the latent code by abstractness and parameterising the factorised posterior distribution with increasingly expressive functions, the VLAE can be trained with the simple ELBO formulation used in non-hierarchical VAEs.

For VaLPACa, this means we separate the shallower-level inference and the deeper-level inference into parallel pathways for inference and generation (1B, orange arrow: *h*_1_ → *h*_2_). We subsequently recombine these levels in the generative network (1B, pink arrow: *f*_*t*_ → *u*_1,*t*_). This approach has two major advantages: 1) It solves the problem of latent variable collapse in which all latent features are loaded onto the lowest-order variables in the hierarchy. 2) It retains a very simple formulation for the variational/evidence lower bound (ELBO) that is easily reparameterisable ^18,23^. We will now outline how VaLPACa does this more explicitly by specifying the generative process, inference model, and cost function.

## VALPACA PROBABILISTIC MODEL

First, we will briefly review the model of LFADS ^2^ and the extensions that make it possible to infer dynamic latent variables from calcium fluorescence traces.

### 3.1 LFADS Generative Model

LFADS assumes that neural activity **x** is generated from a low dimensional non-linear dynamical system with unknown initial conditions subjected to unknown control inputs. The state of the low-dimensional non-linear dynamical system is modelled by the hidden state **g**_**t**_ of a Gated Recurrent Unit (GRU) receiving control inputs *u*_*t*_. The prior for the initial conditions **g**_**0**_ is assumed to be normal with mean *µ*_*g*0_ and variance 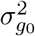. External inputs are assumed to have an autoregressive process prior with process mean *µ*_*u*_, process variance 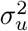, and time constant *τ*_*u*_,

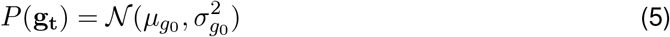

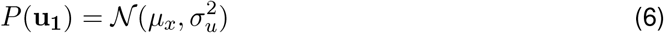

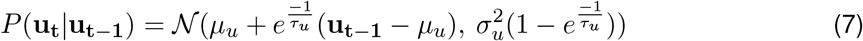

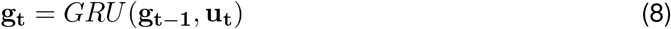

The hidden states of the GRU are then projected to a low-dimensional set of factors *f*_*t*_

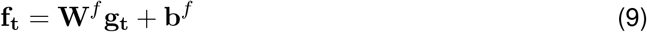

In Pandarinath et al. ^2^, the neural activity being modelled was single-trial spikes extracted from electrophysiological data in macaque and human motor cortex. LFADS assumes a simple observation model for spiking data where the spike count observed in a given time-bin *x*_*t*_ follows an independent Poisson process parameterised by the instantaneous firing rate *λ*_*t*_. To obtain the instantaneous firing rate, low-dimensional factors *f*_*t*_ are linearly projected to the dimensionality of the data and a non-linearity enforcing positivity is applied, in this case the exponential function.

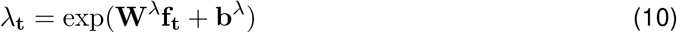

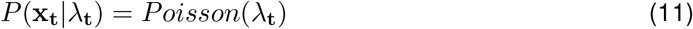

### 3.2 Calcium Fluorescence Generative Model

Before describing how the low dimensional dynamic factors are used to generate calcium fluorescence data, we first describe our probabilistic model for calcium fluorescence transients. We assume that calcium fluorescence transients are generated by an autoregressive process *c*_*t*_ occluded by white emissions noise *ϵ* and perturbed by an unknown spike train **s**_**t**_.

Probabilistic models of unknown spike counts in calcium fluorescence data typically assume a Bernoulli distribution, or some continuous approximation thereof ^12^. This is somewhat problematic given that fluorescence is sampled at a rate of 10-30 Hz, leaving time-bins large enough to observe more than one spike per time-bin. As such, we wanted to use a simple continuous approximation to a Poisson distribution to infer spike counts in each bin. For this purpose, we used a log-normal approximation,

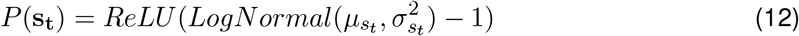

This approximation is easily reparameterizable since we can sample from a lognormal distribution by transforming samples from a normal distribution with an exponential function. The ReLU function and subtraction of 1 from samples ensures that many counts are zero. Furthermore, this parameterization permits a simple closed-form ELBO calculation.

Spike counts are then used to perturb an autoregressive process modelling the slow decay of calcium transients *c*(*t*) after a spike. A simple autoregressive model for calcium transients is the AR(1) process. With discretised time, this can be modelled with Euler updates

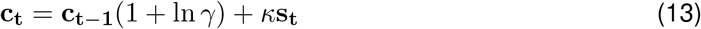

where 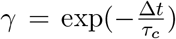, with ∆*t* the size of time bin and *τ*_*c*_ the decay of calcium transients, and with ∆*t* < *τ*_*c*_. This parameterization was chosen to reduce variability during gradient descent, and required restricting the domain *γ* ∈ (0, 1). *κ* is a constant gain term controlling the effect of spikes on fluorescence.

To convert calcium transients to fluorescence *F*_*t*_ we assume a linear independent Gaussian noise model of emissions with constant variance 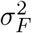,

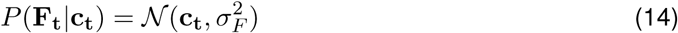

### 3.3 VaLPACa generative model

To tie LFADS and our observation model for calcium fluorescence together, we will adjust the notation slightly to make the hierarchical positioning of latent variables clearer. The variables of the deeper-level model of LFADS are denoted by the subscript *l* = 2, whereas the variables of the shallower-level model of observed calcium fluorescence are denoted by the subscript *l* = 1. For consistency and notation simplicity, we denote variables that are the state of a dynamical system by *g*_*l,t*_ and inputs to the dynamical system as *u*_*l,t*_.

We observe neural activity **x** in the form of single-trial calcium fluorescence traces. We assume that fluorescence follows a simple autoregressive process **g**_**l**=**1**_ perturbed by an unknown set of spike counts **u**_**l**=**1**_ and subject to additive white emissions noise. We also assume that spike counts **u**_**l**=**1**_ are influenced by the state of a nonlinear dynamical system with unknown initial conditions **g**_**l**=**2**,**t**=**0**_ and unknown inputs **u**_**l**=**2**_.

As in LFADS, the nonlinear dynamical system is modelled by a GRU. The hidden state of the GRU *g*_*l* =2,*t*_ is projected to a low-dimensional set of factors **f**_**t**_

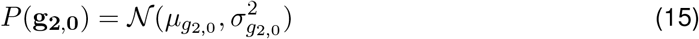

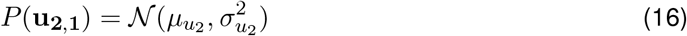

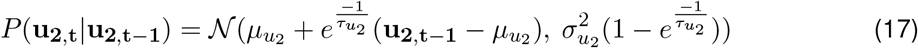

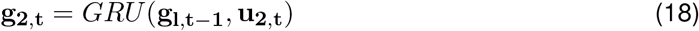

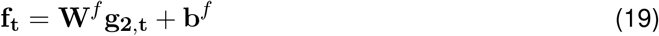

For approximate spike counts to be informed by the dynamic factors **f**_**t**_, we sample according to

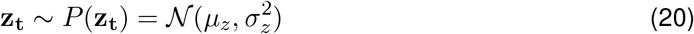

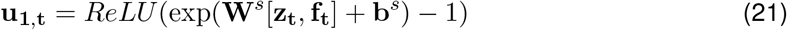

using the continuous approximation to spike counts described in section 3.2.

Although this factorization separates the generation of approximate spikes counts from the dynamic latent factors, for the formulation of the loss function we treat **u**_**1**,**t**_ as if it is generated from a homogeneous Poisson process with intensity *λ*_**t**_ = exp(**W**^*λ*^**f**_**t**_ + **b**^*λ*^), i.e.,

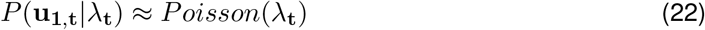

We then use these approximated spikes counts to perturb the state *g*_*l*=1,*t*_ of an AR(1) process

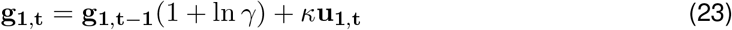

The likelihood of the observed fluorescence in a given time bin **x**_**t**_ is then modelled as normal random variable with a mean dependent on the state of the AR(1) process.

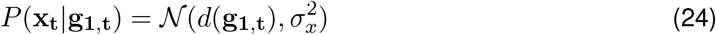

where again *d*(·) is an optional function modelling dye kinetics. In our case *d*(**g**_**1**,**t**_) = **g**_**1**,**t**_.

### 3.4 VaLPACa inference network

The inference network of VaLPACa parameterises the approximate posterior distributions *Q*(*g*_2,0_|*x*), *Q*(*u*_2,*t*_|*x*) and *Q*(*z*|*x*). As per the ladder architecture ^18^ these posterior distributions take the form

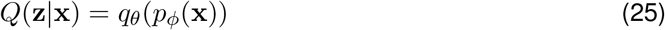

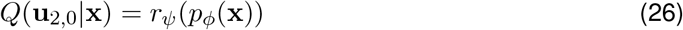

where *p*(·), *q*(·), *r*(·) are neural networks parameterised by *φ, θ, ψ* respectively. The combination of latent space factorization and parameter sharing ensures that dynamics can be disentangled effectively ^18^.

As in LFADS, the approximate posterior distribution over the initial conditions to the nonlinear dynamical system *Q*(*g*_2,0_) is parameterised separately using a bidirectional Gated Recurrent Unit (BiGRU)

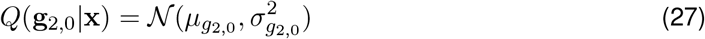

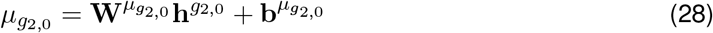

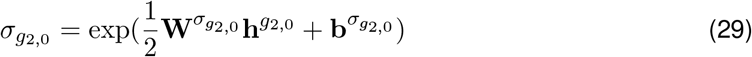

where *h*^*g*^2,0 is the final hidden state of a bidirectional Gated Recurrent unit

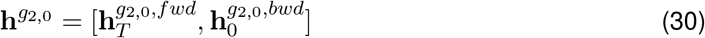

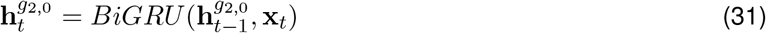

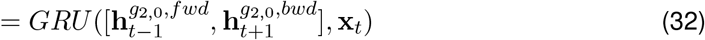

where 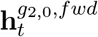 is the state of a GRU running forward sequentially over the input, and 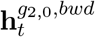 is the state of a GRU running backward sequentially over the data.

The functions *f*_*φ*_(·), *g*_*θ*_(·), *h*_*ψ*_(·) are modelled by pairs of recurrent neural networks. As in LFADS, we use bidirectional GRUs followed by unidirectional controller GRUs receiving representations from the generative network. The bidirectional GRU is defined as before

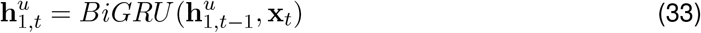

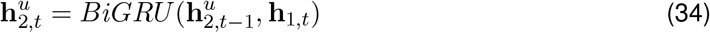

From this point the inference model splits into the two levels of the hierarchy. The hidden state 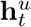 is passed through the a controller GRU which takes feedback from the generated samples

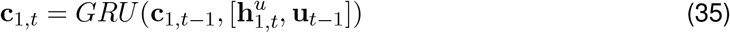

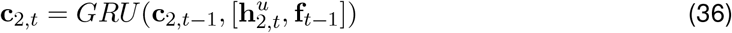

The states of the controller GRU are then linearly transformed onto the parameters of a normal distribution

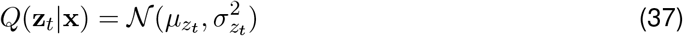

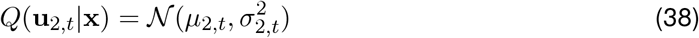

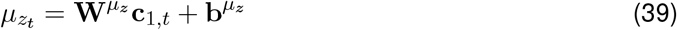

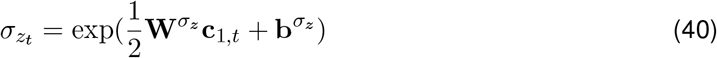

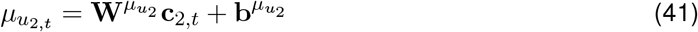

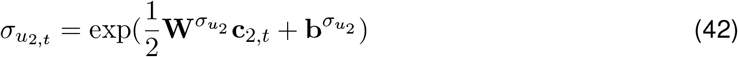

### 3.5 VaLPACa loss function

To construct the cost function of VaLPACa, we start from the evidence lower bound (ELBO)

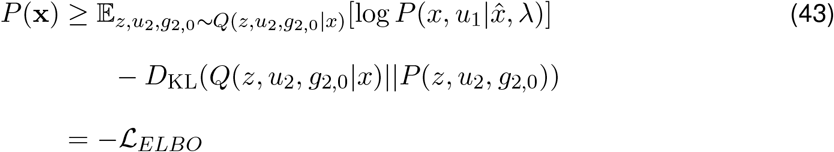

where **x** denotes the observed data and 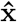 denotes the reconstructed data.

With our model, this factorises as

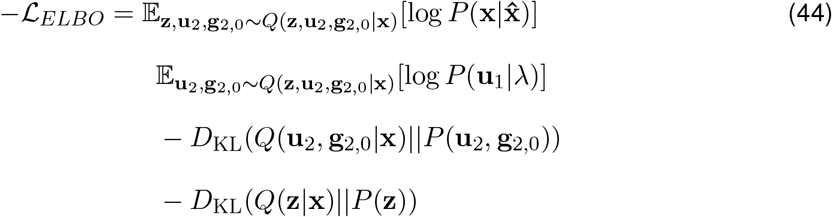

Since *u*_1,*t*_ is a continuous random variable that we are treating as if it came from an independent, homogeneous Poisson process with parameter *λ*_*t*_, we approximate the log-likelihood with

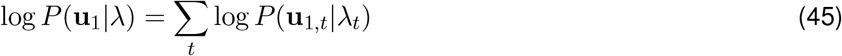

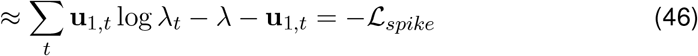

This approximation is very similar to the true log-likelihood of a Poisson distribution. The key difference is that the log-factorial term over the observations is replaced with the identity over the observations, i.e., log *u*_1,*t*_! ≈ *u*_1,*t*_. In deep learning applications, the log-factorial is often dropped entirely from the Poisson log-likelihood (see e.g., the Pytorch implementation and default setting^1^). It is sometimes replaced with a shifted log-gamma function log *k*! ≈ log Γ(*k* + 1) or an approximation via Stirling’s formula 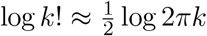. Since Stirling’s formula blows up near zero, and since the log-gamma function has a local minimum between zero and one, they were not deemed appropriate for our model.

However, we observe that this term helps to enforce sparsity in the observations. Since this is desirable for inferred spike counts, we replace this term with what is essentially the *L*_1_ norm of *u*_1,*t*_, since by construction *u*_1,*t*_ is forced to be positive.

The remaining terms of the ELBO were not approximated in any special way. As such we breakdown *L*_*ELBO*_ as follows,

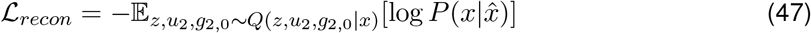

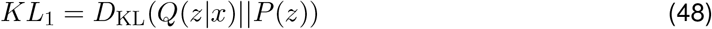

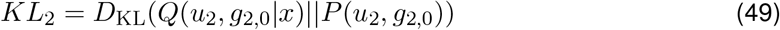

Two additional terms are also added to the cost function, L2 regularizers on the weights of the controller and generator network, and a regularization term for *γ* that both enforces the domain *γ* and allows for incorporation of prior knowledge about likely values of parameter since it is equivalent to the log probability of a Beta distribution over *γ*.

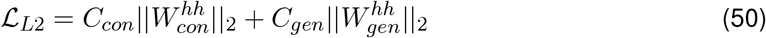

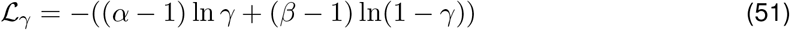

The final cost function of VaLPACa becomes

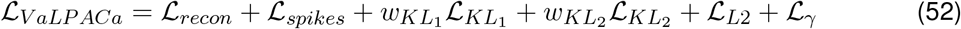

where 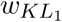 and 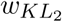 are KL warmup terms that are typically used in training variational autoencoders. These terms slowly transition the model from a traditional autoencoder to a variational autoencoder, which helps to prevent pathological behaviour where the KL terms are minimised too early in training. This typically results in the model then generating trivial solutions. In our case, we use a staggered KL warm-up in which 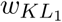 linearly increases from 0 to 1 over epochs 0-100, and then 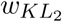 linearly increases from 0 to 1 over epochs 100-200. This allows the model to converge to a solution that overfits spikes to the fluorescence data first, then gradually regularises this proposed set of spikes by ensuring they are well explained by the low-dimensional latent factors.

### 3.6 Training

Parameters of VaLPACa were optimised with ADAM ^24^, with an initial learning rate of 0.01, and batch sizes set at 1/16th of the size of the training data. Learning rates were decreased by a factor of 0.95 when plateaus in training error were detected. As in LFADS training, KL and L2 terms in the cost function were ‘warmed up’, i.e., had a weight *w*_*l*∈(1,2)_ between 0 and 1 applied that gradually increased ^23,25^. Warm-up for the deeper parameters (*l* = 2, blue modules in Figure 1) was delayed until warm-up for shallower parameters (*l* = 1, red modules in Figure 1) was completed. We found in preliminary tests that this delayed warm-up sequence was necessary for better hierarchical inference compared to simultaneous warm-up. As with aggressive inference network training methods ^26^, this prevents issues with training of the deeper-level dynamical system parameters converging more slowly than the shallower-level dynamical system parameters in the initial stages of training.

### 3.7 Implementation and hyperparameters

VaLPACa was implemented in PyTorch 1.7. The majority of experiments were run on a Lenovo Thinkpad P51 with a built-in NVIDIA M1200 GPU. Model hyperparameters for different datasets are shown in table S1 and follow those chosen for LFADS ^2^.

## RESULTS

### 4.1 Lorenz Attractor Synthetic data

#### 4.1.1 Data Description

As an initial test of VaLPACa, we examined its ability to infer the hierarchical dynamical systems of synthetic calcium fluorescence data generated from a simple model of spiking neurons. We examined synthetic data generated from a network with a Lorenz attractor embedded in its dynamics. This tests our ability to recover a ground-truth deeper-level dynamical system from the data, and has been used as a benchmark by many others ^2,27–30^.

In this system, network dynamics were used to generate a set of spike rates in simulated neurons. We then used a model of calcium dynamics and emissions noise to transform the spikes into synthetic fluorescence data. We tested VaLPACa on data generated from two models of synthetic calcium dynamics: a simple linear model equivalent to an AR(1) process, and a nonlinear model in which an AR(1) process is transformed by a Hill function nonlinearity. We then added white noise to the resulting traces. This procedure is described in more detail in Appendix B.

#### 4.1.2 Model Comparison

We measured the performance of VaLPACa by computing *R*^2^ goodness-of-fit with the embedded Lorenz attractor state variables. We also measured the ability of VaLPACa to reconstruct other ground-truth variables in the synthetic data (spike counts, spike rates, and fluorescence traces) using *R*^2^, but these were not as critical for assessing model performance.

VaLPACa performance was compared against two baselines. First, we use LFADS with a Gaussian likelihood observation model to account for fluorescence (Gaussian-LFADS). Second, we consider the situation where spike counts are first estimated separately using the OASIS deconvolution algorithm ^11^, a robust, popular, and computationally inexpensive method of spike extraction widely used by systems neuroscientists ^15,16^. LFADS is then used to infer the deeper-level dynamical system and reconstruct the spike rates from the deconvolved spike trains. We refer to this approach as OASIS+LFADS throughout.

Table 1 compares *R*^2^ goodness-of-fit in reconstructing ground-truth dynamic latent variables in held-out validation data. Table 1 shows VaLPACa is able to reconstruct the Lorenz attractor state for synthetic data with either linear or nonlinear synthetic calcium models (see Table 1 caption for significance test results). Indeed, the inclusion of nonlinearities in calcium dynamics does not affect VaLPACa’s ability to reconstruct the Lorenz attractor state as much as it does with OASIS+LFADS. VaLPACa is also better able to reconstruct other network state variables compared to OASIS+LFADS, as can also be seen in Table 1. However, it is worth noting that high accuracy in reconstructing other network state variables does not appear to be necessary for accurate reconstruction of the underlying Lorenz attractor. This indicates that VaLPACa is better suited to handling uncertainty due to spike-timing and is capable of separating sources of slow dynamics to reconstruct the embedded latent space.

**Table 1:** Comparison of model performance (*R*^2^ mean+sem) on synthetic Lorenz datasets generated with 15 different seeds. A hyphen indicates that the variable cannot be compared, as the model does not infer it. The top row is italicised as the performance of LFADS on this task is considered the upper limit, having no additional observation noise from fluorescence. Results of paired t-tests for testing significance of differences in performance between OASIS+LFADS and VaLPACa within seeds for a) linear calcium model: Lorenz state - *t*_14_ = 2.85, *p* = 0.013; Spike Counts - *t*_14_ = 20.67, *p* < 0.001; Rates - *t*_14_ = 4.81, *p* < 0.001; Fluorescence - *t*_14_ = 16.2, *p* < 0.001. b) nonlinear calcium model: Lorenz state - *t*_14_ = 3.29, *p* = 0.005; Spike Counts - *t*_14_ = 17.16, *p* < 0.001; Rates - *t*_14_ = 6.35, *p* < 0.001; Fluorescence - *t*_14_ = 11.16, *p* < 0.001.

| Calcium | Model | Lorenz state | Spike Counts | Firing Rates | Fluorescence |
| --- | --- | --- | --- | --- | --- |
| NA | *LFADS* | *0.975 ± 0.001* | - | *0.956 ± 0.003* | - |
| Linear | Gaussian-LFADS | 0.818 ± 0.005 | - | 0.701 ± 0.004 | - |
|  | OASIS+LFADS | 0.960 ± 0.001 | 0.580 ± 0.006 | 0.912 ± 0.002 | 0.807 ± 0.005 |
|  | VaLPACa | **0.965 ± 0.001** | **0.817 ± 0.007** | **0.937 ± 0.002** | **0.909 ± 0.003** |
| Nonlinear | Gaussian-LFADS | 0.808 ± 0.005 | - | 0.650 ± 0.006 | - |
|  | OASIS+LFADS | 0.877 ± 0.003 | 0.351 ± 0.004 | 0.682 ± 0.005 | 0.843 ± 0.004 |
|  | VaLPACa | **0.902 ± 0.006** | **0.561 ± 0.010** | **0.756 ± 0.008** | **0.932 ± 0.005** |

For comparison we show example outputs of VaLPACa and OASIS+LFADS in Figure 2. Figure 2A-H provides example reconstructions from VaLPACa and OASIS+LFADS of the fluorescence traces (Fig 2A,E), spike rates (Fig 2B,F), spike counts (Fig 2C,G) and Lorenz attractor states (Fig 2D,H) when using a linear model of synthetic calcium dynamics. Visually, it can be seen that both VaLPACa and OASIS+LFADS achieve a very close fit to the fluorescence traces, spike rates, and Lorenz dynamics. Both models also capture spike-timing, although spike-trains inferred by VaLPACa appear more smoothed due to VaLPACa’s continuous approximation and uncertainty about precise spike-timing. Similarly, both VaLPACa and OASIS+LFADS obtain close reconstructions for the Lorenz attractor states (Fig 2L,P) when a nonlinear model is used to generate synthetic data, although the reconstruction of other network variables 2I-K,M-O) has deteriorated to a much greater extent.

**Figure 2:**
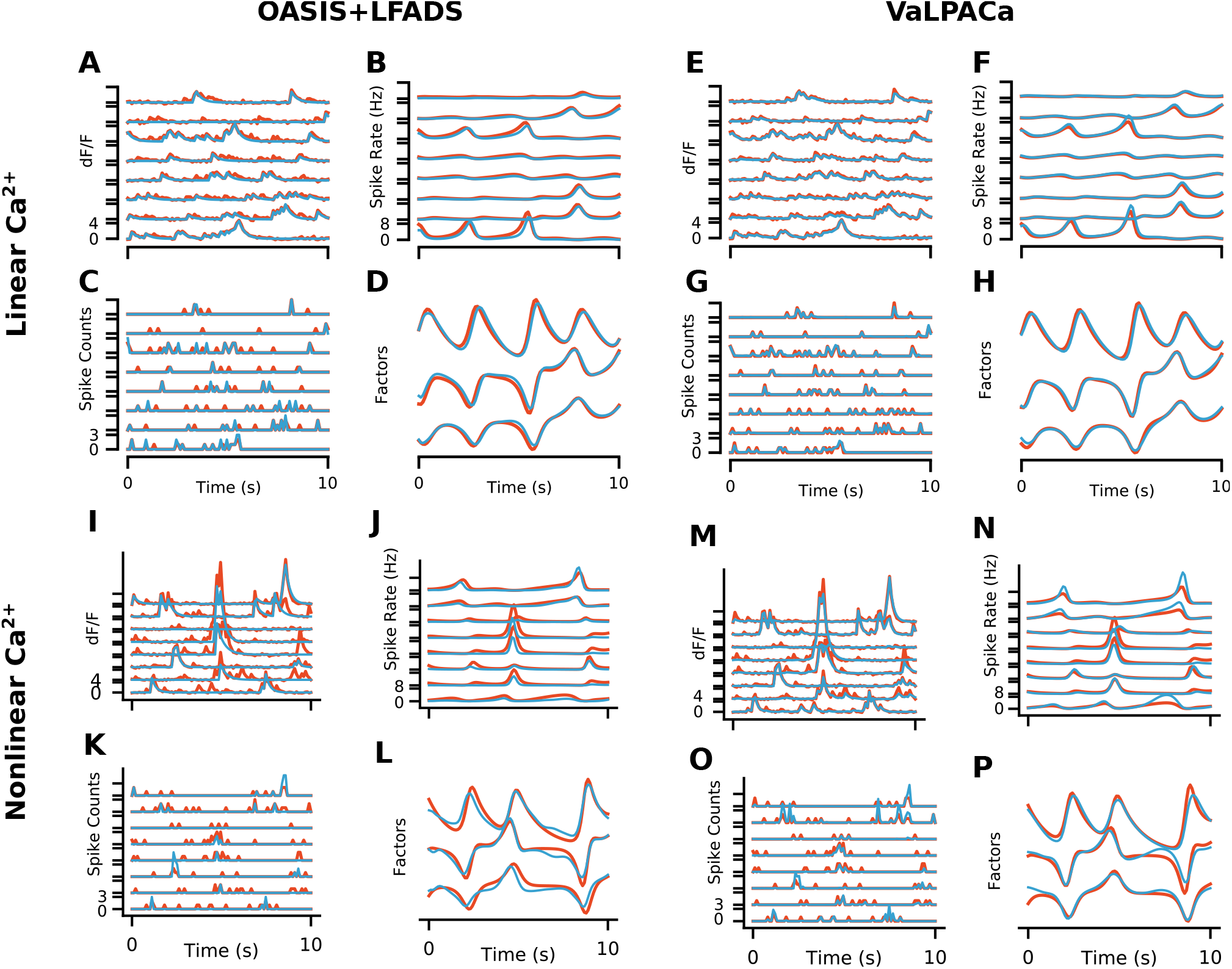
Examples traces of inferred variables compared with ground truth in OASIS+LFADS and VaLPACa for Lorenz attractor synthetic data with either a linear or nonlinear calcium transient generation process. Red: Ground truth, Blue: Reconstruction. A-D) OASIS+LFADS example output for a linear calcium transient generation process for A) Observed fluorescence, B) Spike rates, C) Spike Counts, D) Dynamic factors. E-H) VaLPACa example output for a linear calcium transient generation process for E) Observed fluorescence, F) Spike rates, G) Spike Counts, H) Dynamic factors. I-L) OASIS+LFADS example output for a nonlinear calcium transient generation process for I) Observed fluorescence, J) Spike rates, K) Spike Counts, L) Dynamic factors. M-P) VaLPACa example output for a nonlinear calcium transient generation process for M) Observed fluorescence, N) Spike rates, O) Spike Counts, P) Dynamic factors

Note that VaLPACa reconstructs the Lorenz dynamics almost as well as LFADS does when applied to the spiking data. Of course, it is to be expected that LFADS applied to true spiking data performs better than VaLPACa, since there is an additional source of observation noise from the generation of fluorescence transients. But, the fact that we can get very close to the same level of performance indicates that VaLPACa is effective at performing the same inference on calcium data that LFADS performs on spiking data. We speculate that the uncertainty over AR1 process parameters is overcome in VaLPACa by constraining the reconstructed spike counts with the latent dynamics to make the inference of the observed dynamics process “population-aware”, something which cannot be done when using LFADS applied to spike counts obtained by OASIS deconvolution. This may mitigate errors in spike count reconstruction that occur by deconvolution due to the absence of information about whole-network dynamics. Thus, VaLPACa can use population-level dynamics when conducting the inference of network variables, and this helps it to separate out the shallower-level dynamics more accurately.

### 4.2 Rotational Dynamics in Monkey Motor Cortex

Next, we wanted to test VaLPACa on a real neural dataset to which LFADS has previously been applied, and successfully used to uncover meaningful latent dynamics. Specifically, it has been shown previously that rotational dynamics underlie neuronal responses in monkey and human motor cortex during reaching behaviour ^1^. LFADS has been successful in uncovering these known rotational dynamics for single-trial spikes recorded from primary motor cortex (M1) and dorsal premotor cortex (PMd) in macaques ^2^. To test whether VaLPACa could do the same, we took the original spiking data and converted it into semi-synthetic calcium traces, using the same model of calcium dynamics that was used for synthetic data generation in the previous sections (Fig 3A). We used data from the monkey electrophysiology dataset previously described, along with the reaching task, by Churchland et al. ^1^, which was kindly provided to us by the original authors. Briefly, monkeys were trained to reach a target under 108 different reach conditions while multi-electrode recordings were made in M1 and PMd. Reaches started from a specified location on a screen, and monkeys were rewarded for correctly reaching toward a target while avoiding on-screen obstacles (Fig 3B).

**Figure 3:**
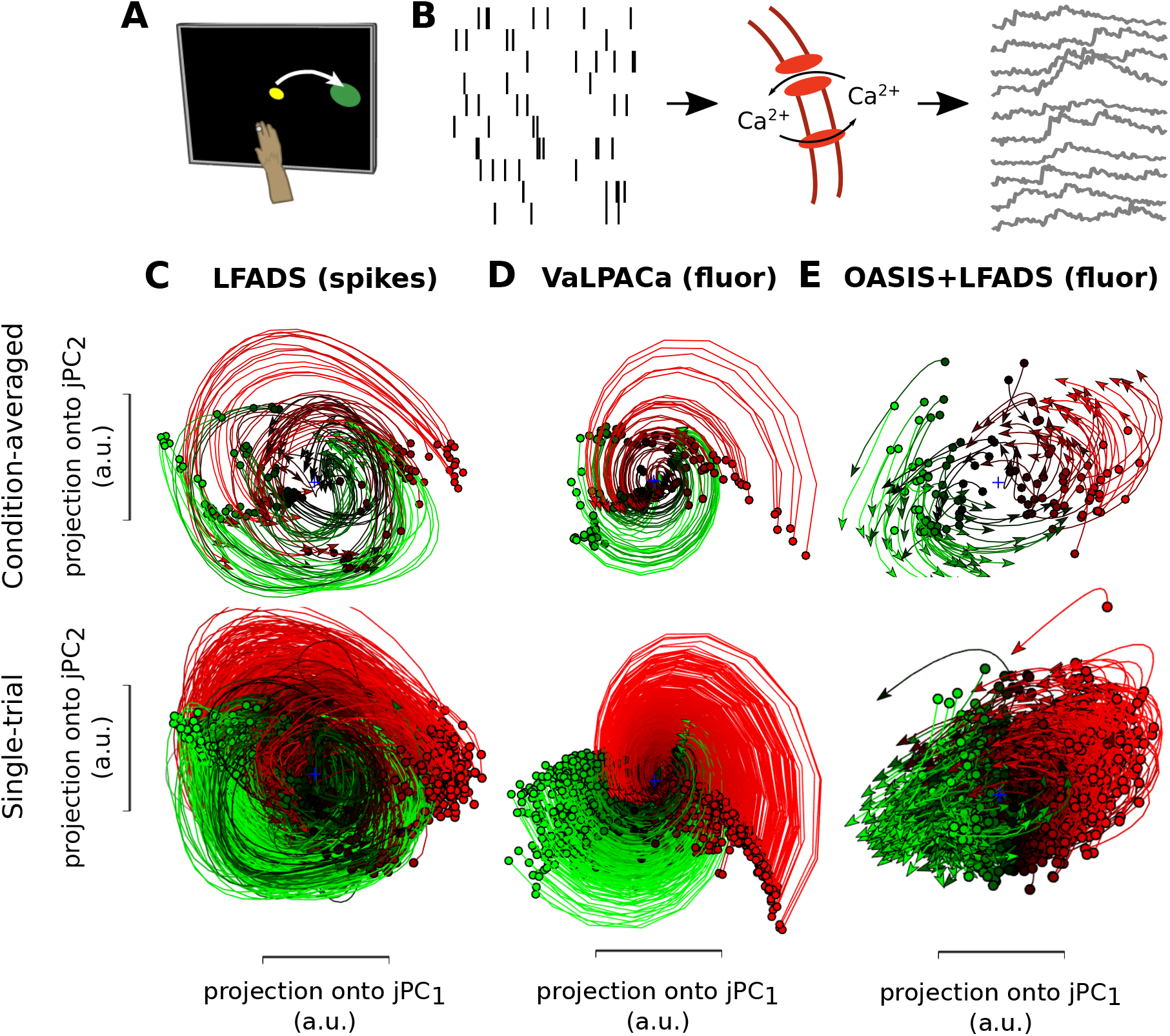
A) Schematic for the process of converting spikes to calcium traces, and B) for the reaching task. C) Rotational dynamics inferred from condition-averaged (Top) and single-trial spikes (Bottom) using LFADS. D) Same as C), but for VaLPACa. E) Same as C), but for OASIS+LFADS. Traces are coloured based on their initial state values along jPC1 (from green to red for increasingly large values).

First, to replicate the previous LFADS results, we applied jPCA ^1^ to condition-averaged firing rates (trial averages for each of the 108 reach conditions), as well as single-trial firing rates inferred from spike data using LFADS. jPCA is a dimensionality reduction technique that finds orthogonal projections capturing rotational dynamics that explain variability in firing rates. The original LFADS paper showed that rotational dynamics explained a large amount of the variance in firing rates. As in the original papers, we identified both condition-averaged (Fig 3C – top) and single-trial (Fig 3C – bottom) rotational dynamics from firing rates inferred by LFADS, which explained a large amount of the variance (*R*^2^ = 0.81), thereby replicating the original findings. Then, we evaluated VaLPACa’s ability to uncover the rotational dynamics of both condition-averaged and single trials from firing rates. As shown in Fig 3, our model also successfully uncovers rotational dynamics from calcium traces, explaining a large amount of variance (*R*^2^ = 0.78) for both condition-averaged (Fig 3D – top) and single trials (Fig 3D – bottom). These results demonstrate that VaLPACa is capable of identifying latent dynamics in real neural data, similar to LFADS, even when the spiking data is transformed by calcium dynamics and emissions noise. We also found that the OASIS+LFADS approach was not as successful as VaLPACa in uncovering rotational dynamics, explaining only half the variance (*R*^2^ = 0.53, Fig 3E). This result corroborates the importance of the hierarchical modelling of calcium and neuronal dynamics in VaLPACa.

### 4.3 Unexpected Stimulus Detection in Mouse Primary Visual Cortex

Finally, we wanted to test VaLPACa on an entirely new calcium imaging dataset, to determine whether VaLPACa can infer dynamic computational factors that carry information about relevant features of the outside world. We chose to analyse data from mouse primary visual cortex (VisP), a region widely studied in systems neuroscience research where calcium imaging is a standard tool. There is evidence that the visual cortex of mammals performs a predictive function, anticipating expected stimuli ^31–34^. As a result, unexpected stimuli can induce perturbations in network dynamics ^33,34^. Thus, we wanted to determine whether VaLPACa could infer dynamic computational factors that carry information distinguishing unexpected from expected visual stimuli.

To this end, we trained VaLPACa on calcium imaging data from the mouse visual cortex collected in collaboration with the Allen Institute for Brain Science ^35^ (for a detailed description, see Appendix C). While awake behaving mice were presented with visual stimuli on a screen (Fig 4A), calcium fluorescence responses in cortical layer 2/3 of VisP were recorded using 2-photon microscopy (Fig 4B). The mice were familiarised with sequences of stimulus frames that followed simple probabilistic rules. Briefly, each sequence consisted of four randomised Gabor pattern frames followed by a grey screen (A, B, C, D, grey) with orientations drawn for each trial from the same distribution. After familiarization with this sequence, the expected D frame was replaced in ∼7% of trials by a Gabor pattern (U) with orthogonal orientations (Fig 4C).

Figures S2 show examples of actual fluorescence traces, the corresponding inferred spike counts and spike rates, and inferred latent factors generated as the output of the VaLPACa and OASIS+LFADS models. Notably, we see that the inferred latent factors of VaLPACa are smoother and less noisy compared to those of the OASIS+LFADS.

**Figure 4:**
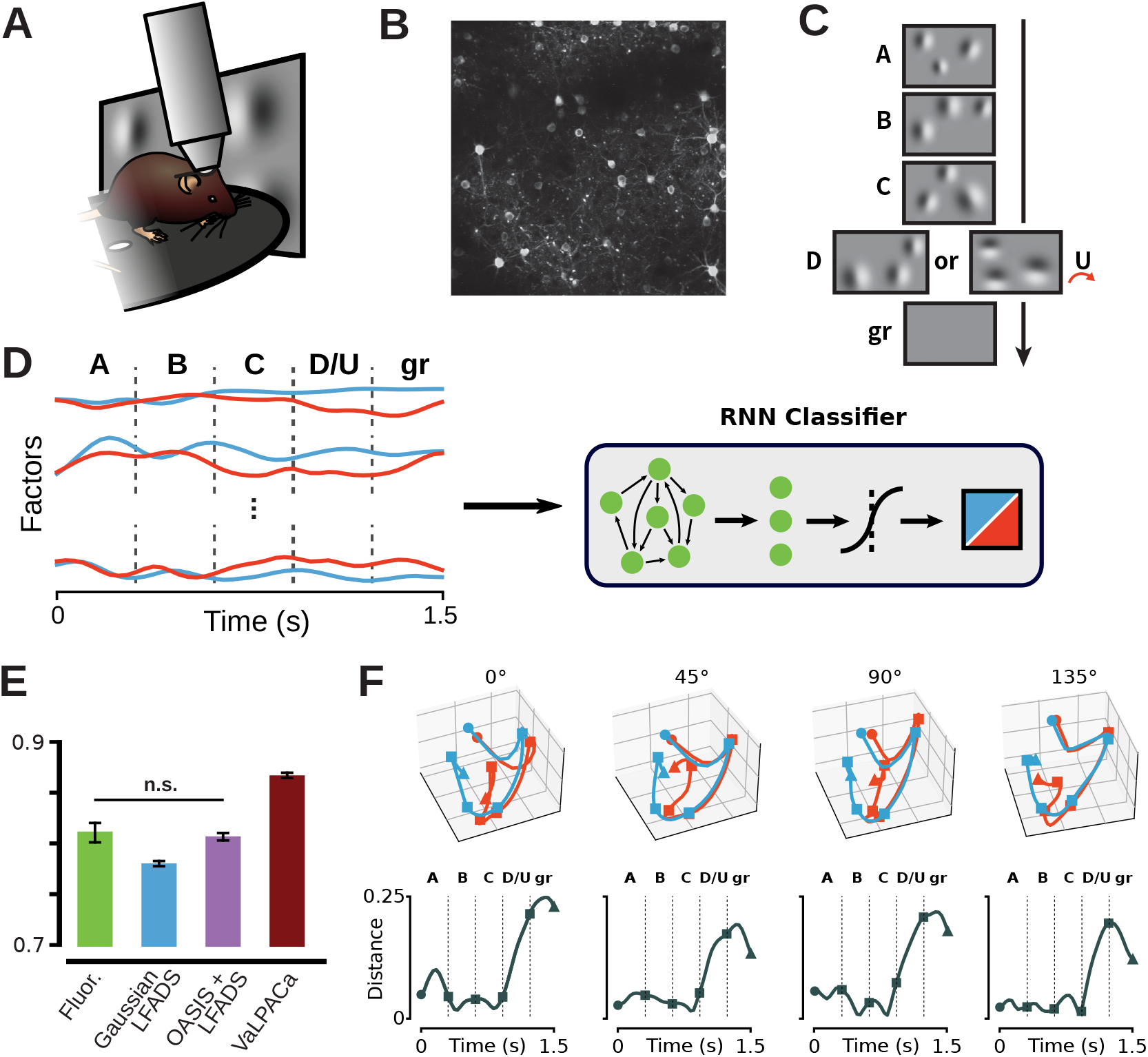
A) Schematic of 2-photon calcium imaging recording setup. Mice are head-fixed on a running wheel under the imaging objective, and visual stimuli are projected onto a screen to the side, (B) 2-photon calcium imaging recording plane, C) Schematic example of visual stimuli for expected (A-B-C-D-grey) and unexpected (A-B-C-U-grey) trials. D) Schema of the non-linear model trained to decode expected vs. unexpected stimulus trials. Example latent factors inferred by VaLPACa are plotted for an expected (blue) and an unexpected (red) trial. Each stimulus frame (A, B, C, D/U, grey) is labelled, and its onset and offset in the trial are marked with dotted lines. Latent factors are passed through GRU and linear modules (green), followed by a sigmoid decision function. E) Average recall (mean ± sem) of expected vs. unexpected trials across non-linear decoders trained on principal components of fluorescence traces (0.815 ± 0.011, green), Gaussian-LFADS factors (0.776 ± 0.004, blue), OASIS+LFADS factors (0.810 ± 0.005, purple), and VaLPACa factors (0.871 ± 0.004, maroon). All pairs but one (fluor. vs. OASIS+LFADS factors, marked with n.s.) were significantly different at p < 0.05 in 2-tailed independent t-tests with Bonferroni correction for multiple comparisons. F) Top: 3D projection of factors in principal component space separated by unexpected (red) and expected (blue) stimulus trials for Gabor frames sequences mean orientations of 0^º^, 45^º^, 90^º^, and 135^º^. Circle = trial start, square = frame change, triangle = trial end. Bottom: Euclidean distance between unexpected and expected trial factor trajectories.

To determine if salient information about these unexpected stimulus features was present in the inferred factors from VaLPACa, we trained a non-linear decoder to classify whether the inferred factors came from a trial with an expected D frame or an unexpected E frame (Fig 4 for details, see Appendix C). We also trained the non-linear decoder on factors inferred by OASIS+LFADS, and on fluorescence traces reduced in dimensionality with principal components analysis. To compare the models fairly, despite the imbalance between expected and unexpected trial frequencies, we measured average recall. We find that using the VaLPACa factors leads to the highest performance on stimulus-trial identity decoding, with the average recall being significantly higher than when OASIS+LFADS factors are used (Fig 4E). This indicates that VaLPACa infers latent dynamics from real visual cortex calcium imaging data that reflect the predictability of visual stimuli, corroborating the ability of VaLPACa to infer meaningful dynamic factors in real data.

To examine why discrimination performance improved for latent factors inferred from VaLPACa, we visualised factor trajectories in a 3D space using PCA, and compared the point-wise distance between trajectories (Fig 4F, Fig S3). The distance between trajectories diverges starting with the surprise frame (Fig 4F), and furthermore, the divergence in trajectories is still present when we compare trials with matched orientations during the surprise frames (Fig S3). This clearly indicates that the factors are learning to represent surprise information separate from orientation information. This corroborates the findings of ^35^, who demonstrated that L2/3 pyramidal cells receive top down input representing surprise information.

## DISCUSSION

In this paper we presented VaLPACa, a hierarchical recurrent variational autoencoder model capable of reconstructing latent computational dynamics. We confirmed VaLPACa’s ability to reconstruct known underlying dynamics using synthetic datasets where ground-truth was known. We also showed that VaLPACa is able to infer sensorimotor dynamics from real neural recordings. This indicates that VaLPACa is a promising method for analyzing calcium imaging data in neuroscience.

There are two key advantages of VaLPACa over the use of deconvolution of calcium traces followed by application of LFADS. The first is that we can obtain measures of the uncertainty in both the latent dynamics, and the latent spike counts. The second is that VaLPACa performs better than OASIS+LFADS when nonlinearities in calcium dynamics are present, as is the case in real experimental data where there are many sources of calcium influx related to spikes.

While we have demonstrated how powerful VaLPACa can be as a tool for analysing calcium imaging data, it is likely not the only way to approach this problem. Alternative continious approximations to spike counts could still be explored, such as the zero-inflated Gamma model ^36^ or techniques using the Gumbel-Softmax trick ^20,37^. An interesting approach that we did not explore is to generate fluorescence traces directly from firing rates by marginalising over spike counts ^38^, in which case a ladder architecture might not be necessary. Additionally, while we have shown that VaLPACa has better performance than a Gaussian-LFADS model on 2-photon imaging data, a simple Gaussian observation model will be effective for inferring nonlinear latent dynamics in wide-field calcium imaging data where spiking noise from individual cells does not dominate the signal so heavily, as shown by ^39^. Finally, while we found that a linear model of calcium dynamics was sufficient to infer nonlinear latent dynamics, it would be a relatively simple extension to explore nonlinear calcium dynamics models.

We designed VaLPACa to fit into a much broader class of artificial neural network models, namely *sequential variational ladder autoencoders*, in which different layers of recurrent neural networks can be used to infer and generate different layers of a hierarchical dynamical system. In this sense, this class of models is modular and composable, meaning it could readily be adapted to other domains. For example, it should be possible to replace LFADS as the deeper-level dynamical system model with any other differentiable model. Likewise the same is true for our AR1 based model of calcium dynamics: we could replace the calcium dynamics model with other models as required by the experimental set-up. Furthermore, there is no need to stop at two layers in the hierarchy; the brain is comprised of many interconnected recurrent neural networks sending long range signals to one another, and it should be possible to add in additional modules to capture additional hierarchies and dynamics within the brain. To date, we are unaware of any other instances of models in this class, however we believe our results from VaLPACa demonstrate that sequential variational ladder autoencoders are a useful model class for developing deep hierarchical inference algorithms.

In summary, VaLPACa is a new, open source tool for inferring hierarchical dynamics from calcium imaging data that also has great potential for being modified and applied to other data modalities. This could be of real benefit to the thousands of neuroscience laboratories around the world conducting calcium imaging experiments.

## Author Contributions

LYP, BAR designed model architecture. LYP, SB implemented model architecture. LYP, SB, CJG prepared and analyzed datasets. LYP, SB, CJG, BAR wrote the paper.

## Acknowledgments

This research was supported by an NSERC Discovery Grant to BAR (RGPIN-2014-04947), a CIFAR Catalyst grant to BAR, the CIFAR Learning in Machines and Brains Program, and a Postdoctoral Scholarship to LYP from the Canadian Open Neuroscience Platform. Thanks to Mark Churchland, Matt Kaufman and Krishna V Shenoy for providing Macaque motor cortex data used in this the paper. KVS received funding support from NIH Director’s Pioneer Award (1DP1OD006409), NIH NINDS EUREKA Award (R01-NS066311), NIH NINDS CRCNS (R01-NS054283) and DARPA-DSO REPAIR (N66001-10-C-2010). The 2-photon imaging data presented herein were obtained at the Allen Brain Observatory as part of the OpenScope project, which is operated by the Allen Institute for Brain Science. We thank Jérôme Lecoq, Jed Perkins, Sam Seid, Carol Thompson, and Ryan Valenza for work regarding collecting and managing the 2-photon imaging data. This work was supported by the Allen Institute and in part by the Falconwood Foundation. We thank Allan Jones for providing the critical environment that enabled this large-scale team effort. We thank the Allen Institute founder, Paul G Allen, for his vision, encouragement, and support. And we thank Christof Koch for his vision for the OpenScope project.

## VARIABLE GLOSSARY

**Table S1:** Glossary of variables, with descriptions and dimensionality across datasetsss

| Variable | Description | Dimensions |  |  |
| --- | --- | --- | --- | --- |
|  |  | Lorenz | M1/PMd | VisP |
| $x_t$ | Fluorescence signal | 30 | 202 | 96 |
| $h_{1,t}$ | Calcium dynamics embedding encoder state | 128 | 200 | 128 |
| $h_{2,t}$ | Computational dynamics embedding encoder state | 64 | 100 | 128 |
| $c_{1,t}$ | Calcium dynamics embedding controller state | 128 | 200 | 128 |
| $c_{2,t}$ | Computational dynamics embedding controller state | 0 | 0 | 64 |
| $g_{1,t}$ | Calcium dynamics embedding state | 100 | 200 | 128 |
| $z_{2,t}$ | Computational dynamics embedding state | 64 | 100 | 200 |
| $u_{1,t}$ | Approximate spike counts | 30 | 202 | 96 |
| $u_{2,t}$ | Computational dynamics perturbations | 0 | 0 | 1 |
| $g_{1,t}$ | Calcium dynamics | 30 | 202 | 96 |
| $f_t$ | Dynamic factors | 3 | 40 | 32 |
| $\hat{x}_t$ | Reconstructed fluorescence signal | 30 | 202 | 96 |

## SYNTHETIC DATA GENERATION

As discussed in the main text, synthetic calcium fluorescence data was generated by embedding a Lorenz Attractor in the dynamics of a spiking neural network. Two models of calcium dynamics were then used to transform spikes to calcium transients, with additive emissions noise added to the resulting traces.

The Lorenz Attractor is a nonlinear dynamical system with 3 states *x*_1_, *x*_2_, *x*_3_ commonly used to study chaotic dynamics. The dynamical system is defined by,

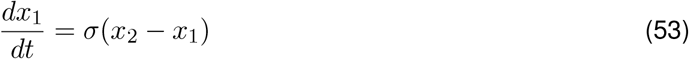

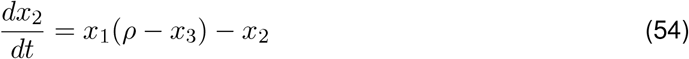

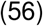

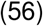

This system was parameterised in its typical chaotic regime with *σ* = 10, *β* = 8*/*3, *ρ* = 28. These states were then normalised to have a mean of zero and a range of [–1, 1].

The state of this system **x** was then randomly projected onto the firing rate of a population of *N* = 30 neurons,

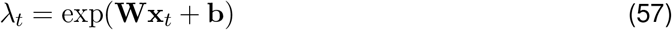

with 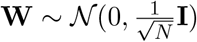, and **b** = **1**. Spike counts in time-bin *t* for neuron i *s*_*i*_ were then sampled using a Poisson distribution

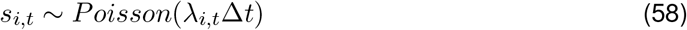

where ∆*t* is the width of a time-bin. For our simulations, we used ∆*t* = 0.1*s*.

Calcium fluorescence traces were then modelled in one of two ways. The first method modelled calcium concentration in neuron i *c*_*i*_ as an exponentially decaying variable with time constant *τ* = 0.3 and perturbations by spikes

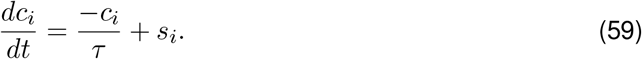

Fluorescence was then modelled by adding white emissions noise with standard deviation *σ*_*F*_ = 0.2.

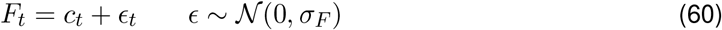

The second method for generating synthetic calcium traces added an intermediary step between calcium influx and observed fluorescence. As previously, spikes were integrated with a slow-varying exponentially decaying variable *c* as described in Equation 59. A hill function was used to capture the nonlinear binding kinetics of calcium to the indicator dye,

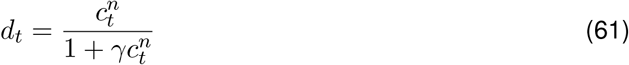

with hill coefficient *n* = 2, and *γ* = 0.0001. These parameters were chosen based on parameter fits in Deneux et al. ^13^. Finally white noise was added to these resulting traces to provide synthetic fluorescence traces,

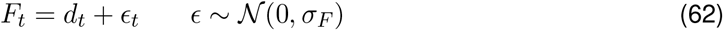

## MOUSE VISP DATASET

The 2-photon calcium imaging data used here is part of the OpenScope project dataset ^35^, collected by the Allen Institute for Brain Science in Seattle, WA. All animal procedures were approved by the Institutional Animal Care and Use Committee at the Allen Institute for Brain Science. Neuronal activity was recorded in head-fixed, awake Cux2-Cre mice on a running wheel ^40^ (Fig 4A) expressing the calcium indicator GCamP6f ^41^ in layer 2/3 pyramidal cells of VisP (Fig 4B). The mice were first habituated to a repeating, expected stimulus over 6 days, after which unexpected trials were introduced. The stimuli were adapted from ^42^. In each expected trial, 4 consecutive sets of 30 Gabor patches appeared in sequence for 300 ms each, followed by 300 ms of grey screen (A, B, C, D, grey). For each set, the locations and sizes of the Gabor patches were held constant within a session. However, within each trial, each Gabor patch’s orientation was sampled from a von Mises distribution, with trial mean sampled from {0, 45, 90, 135}^º^ and standard deviation 0.25^º^. In expected trials, occurring ∼7% of the time, the D set was replaced with a distinct set E, with its own locations and sizes. In addition, the Gabor patch orientations for E sets were sampled from a von Mises distribution with mean shifted positively by 90^º^ (Fig 4C). Processed fluorescence (dF/F) traces were extracted from the calcium imaging recordings for the putative neurons identified, as described in ^35,40^.

**Figure S1:**
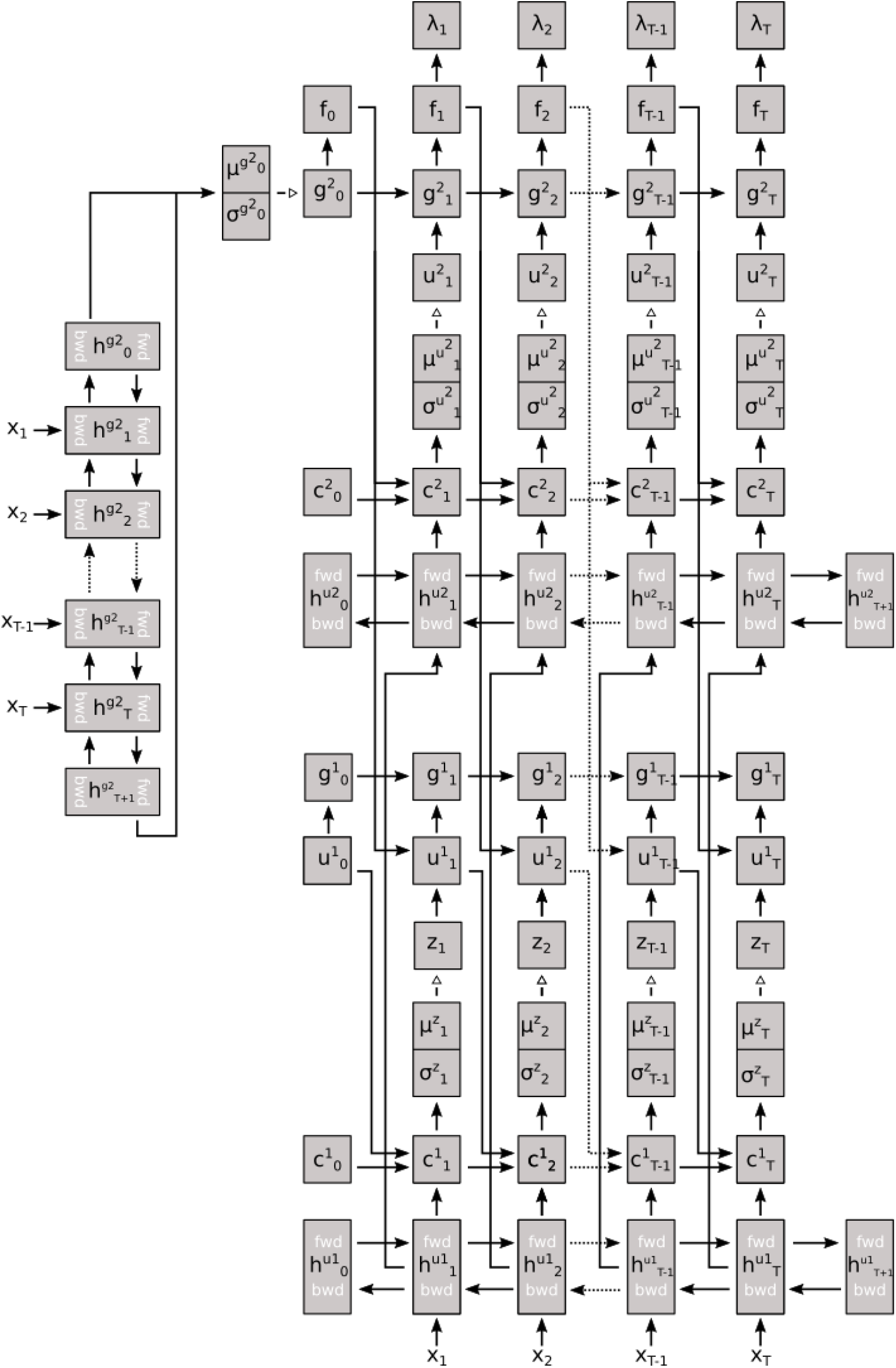
Directed acyclic graph for hierarchical model. Solid arrows denote deterministic mappings, open arrows denote sampling steps.

**Figure S2:**
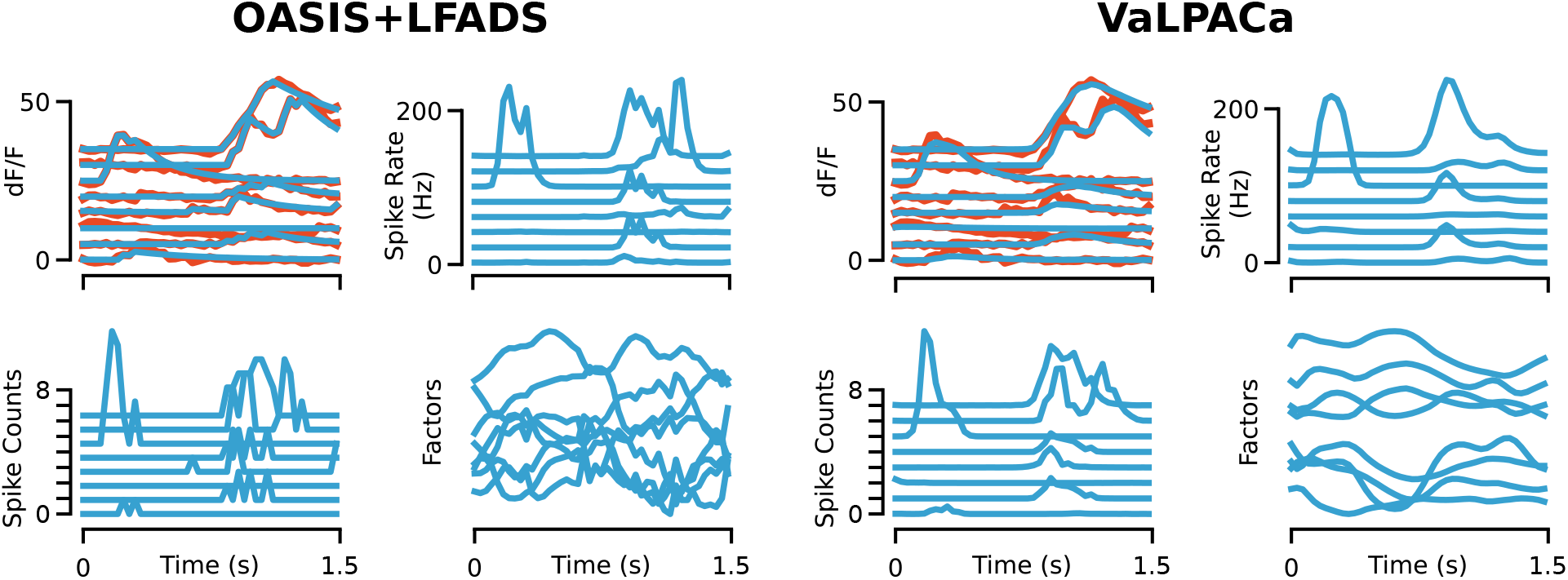
Samples of actual and reconstructed fluorescence traces, inferred spike count, spike rates and latent factors in OASIS+LFADS and VaLPACa

**Figure S3:**
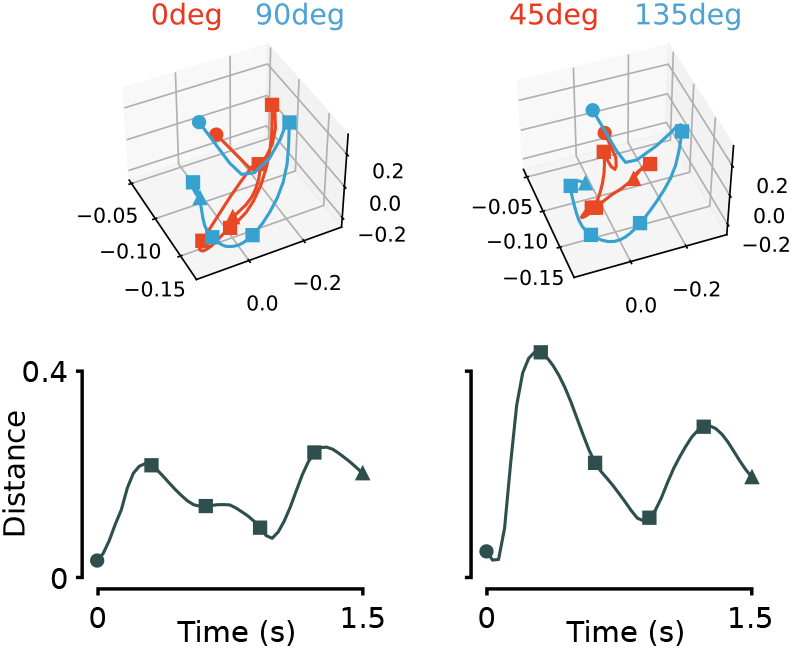
Top: 3D projection of factors in principal component space separated by unexpected (red) and expected (blue) stimulus trials for Gabor frames sequences where the surprise and non-surprise frame have the same mean orientation, and all other frames have orthogonal mean orientations. Circle = trial start, square = frame change, triangle = trial end. Bottom: Euclidean distance between unexpected and expected trial factor trajectories. Two pairs of conditions are omitted as they do not match due to Gabor patch asymmetry.

